# Concatemeric Broccoli reduces mRNA stability, aggregates and induces p-body formation

**DOI:** 10.1101/2020.12.07.414458

**Authors:** Marco R. Rink, Marisa A.P. Baptista, Thomas Hennig, Adam W. Whisnant, Natalia Wolf, Jürgen Seibel, Lars Dölken, Jens B. Bosse

## Abstract

Fluorogenic aptamers are an alternative to established methodology for real-time imaging of RNA transport and dynamics. We developed Broccoli-aptamer concatemers ranging from 4 to 128 substrate-binding site repeats and characterized their behavior fused to an mCherry-coding mRNA in transient transfection, stable expression, and in recombinant cytomegalovirus infection. Concatemerization of substrate-binding sites increased Broccoli fluorescence up to a concatemer length of 16 copies, upon which fluorescence did not increase and mCherry signals declined. This was due to the combined effects of RNA aptamer aggregation, a nuclear export defect and reduced RNA stability. Unfortunately, both cellular and cytomegalovirus genomes were unable to maintain and express high Broccoli concatemer copy numbers, possibly due to recombination events. Overexpression of Broccoli-tagged mRNA led to the formation of p-bodies. However, Broccoli RNAs did not localize to these sites. Interestingly, negative effects of Broccoli concatemers could be partially rescued by introducing linker sequences in between Broccoli repeats warranting further studies. Finally, we show that even though substrate-bound Broccoli is easily photobleached, it can still be utilized in live-cell imaging by adapting a time-lapse imaging protocol.

## Introduction

RNA aptamers are short RNA sequences that exert specific binding abilities to a given biological structure or small molecule. Fluorogenic RNA aptamers in this regard are characterized by their ability to bind a small molecule and greatly enhance its fluorescence potential. This has been first reported with an aptamer called Spinach [1] (the name due to its green fluorescence) and been further expanded over the recent years with other aptamers emerging, such as the RNA Mangos [2], Broccoli [3] or Corn [4]. The most prevalent RNA aptamers are RNA mimics of GFP, termed as such by the very similar peak excitation and emission wavelengths. Mechanistically these aptamers consist of binding a GFP mimicking small molecule fluorophore, thereby inhibiting vibration and isomerization, forcing fluorescence as the only remaining mechanism to dissipate energy [5]. As such, the aptamer itself becomes visible when its substrate is present, the substrate in its unbound state is non- or only lowly fluorescent. For this, the most widely used substrate is DFHBI ((5Z)-5-[(3,5-Difluoro-4-hydroxyphenyl)methylene]-3,5-dihydro-2,3-dimethyl-4H-Imidazol-4-one) or a derivative thereof, which are generally membrane permeable.

Due to these features, RNA aptamers, when fused to an RNA of interest, can be utilized to detect RNAs in living cells by all common methodologies that utilize fluorescence, e.g. flow cytometry or fluorescence microscopy and constitute an alternative to the “gold-standard” for RNA imaging, the MS2-tagging system [6]. Tandem arrays of MS2-tags fused to a given RNA have tremendous RNA tracking potential potentially providing single molecule resolution [7]. An optimal RNA tag would enable researchers to visualize single RNA molecules without perturbing their normal biological properties and behavior. Here, RNA aptamers were proposed to potentially have a smaller impact on the biological properties of the tagged RNA molecule.

Fluorescent RNA aptamers have now been used to elucidate many fundamental RNA biology processes, including transcription rates [4] and RNA aggregate studies [8]. Especially when high concentrations of RNA are present within a structure, the aptamer approach should provide insights into cell’s RNA biology. As such, p-bodies, stress granules, rRNAs and small non-coding RNAs and their dynamics are especially interesting. Single aptamer tags so far have not been successful in the tracking or quantification of single molecules though, but reminiscent of the MS2-tag concatemers, tandem arrays could greatly empower the scope of the methodology. Recently, the tandem array approach has been successfully utilized, in bacteria [9], and, with a new substrate, to visualize overexpressed RFP carrying mRNA molecules in mammalian cells [10]. However, the introduction of multiple extensively structured exogenous sequences into a given RNA molecule may alter the physiological behavior of the respective molecules. This question has been addressed by Xing *et al.* and they determined no effect of the repeated Broccoli RNA aptamer on mRNA stability [10]; quite different to our experience with it.

Here, we report on the development of high copy number Broccoli RNA aptamer tandem arrays and their effects on RNA export, localization and stability, with the scope of using the system to visualize and track viral RNAs in productive virus infection.

## Results

In order to perform high-resolution microscopy for single molecule imaging in mammalian cells, we generated tandem arrays of the aptamer Broccoli. We expanded the number of binding sites for the fluorochrome DFHBI from 4 to 128 copies by repetitive duplications (strategy described in the methods section and in **Fig. 1A**; refer to **Table 2** for nomenclature and insert sequence length). The coding sequence (CDS) of the fluorescent protein mCherry was inserted upstream of the Broccoli tandem arrays to monitor their effects on protein expression.

**Figure 1.**
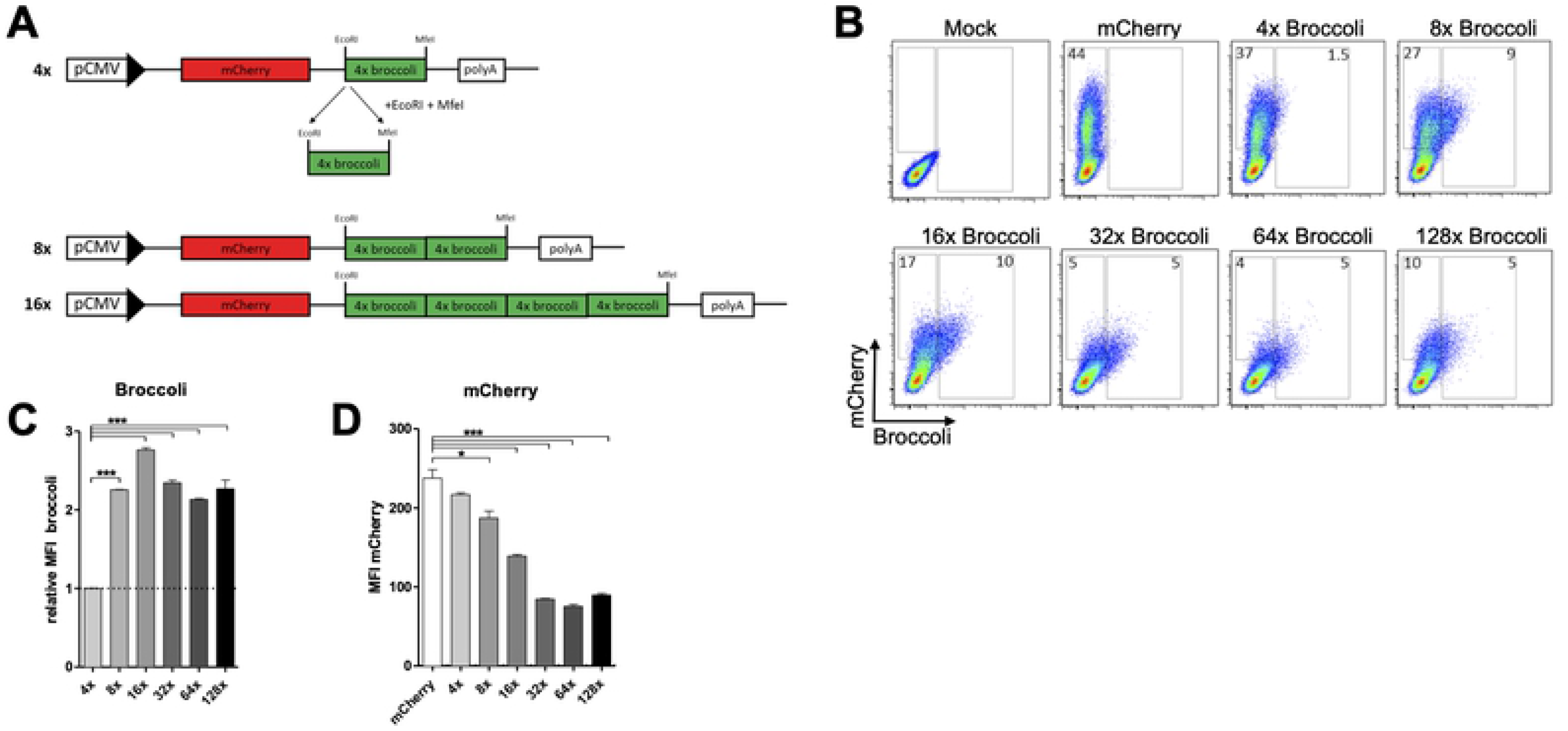
Broccoli tandem array synthesis and expression. (a) Cloning schematic of the constructs containing different copy numbers of Broccoli. (b, c, d) 293T cells were transfected with mCherry or 4×, 8×, 16×, 32×, 64× or 128× Broccoli and analyzed 24 hours post-transfection by flow cytometry. Percentage of Broccoli and mCherry positive cells shown in dot plots (b) and mean fluorescence intensity of Broccoli (c) and mCherry (d) compared between constructs. (c) The mean fluorescence intensity (MFI) of Broccoli in the graph was normalized to the 4× Broccoli construct and the MFI of the remaining constructs was calculated as follows: ⦗ (MFI **X** xBroccoli): (4xBroccoli) ⦘. Data representative of three independent experiments. Statistical analysis performed by t-test (c, d). * P<0.05, **P<0.01, *** P<0.001

To determine the fluorescence intensity of each Broccoli tandem array, 293T cells were transfected with the Broccoli-expressing plasmids and 24 hours later the cells were collected, resuspended in PBS buffer containing DFHBI, and analyzed by flow cytometry (**Fig. 1B**). While 4× Broccoli only resulted in a weak green fluorescence in mCherry positive cells, green fluorescence doubled from 4× to 8× Broccoli. Although a small further increase in green signal was observed for 16× Broccoli, no further increase was observed thereafter (**Fig. 1C**). While 44% of the cells were mCherry-positive when transfected with an mCherry-only control plasmid, the percentage of mCherry-positive cells decreased with increasing size of the Broccoli reporter. This was to be expected and consistent with a decrease in transfection efficiency with increasing plasmid size (**Fig. 2A, Supplementary Fig. 1A**). While only a modest decrease in mCherry expression strength was observed up to 8× Broccoli, mCherry expression dropped markedly with increasing numbers of Broccoli repeats (**Fig. 1D**). Combined, these data indicate that increasing numbers of Broccoli repeats beyond 8× impairs gene expression of the reporter.

**Figure 2.**
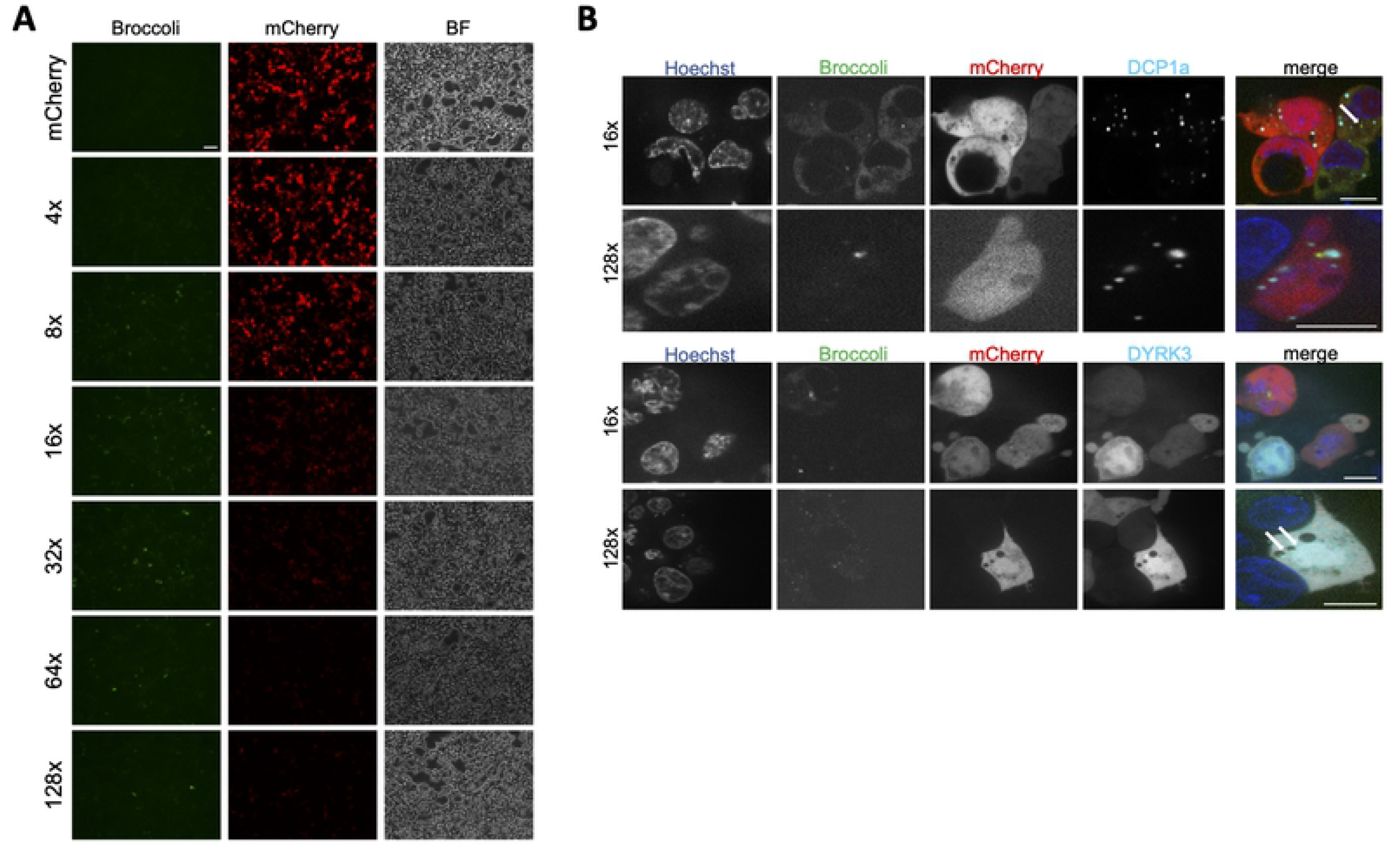
Broccoli exhibits punctate morphology that does not co-localize to p-bodies or stress granule markers. (a) 293T cells were transfected with mCherry, 4×, 8×, 16×, 32×, 64×, 128× Broccoli plasmids and analysed by microscopy for localization and morphology. (Scale bar = 75 μm)(b) To assess the origin of aggregate formation of Broccoli, 293T cells were co-transfected with plasmids expressing DCP1a-SNAP or DYRK3-SNAP to label p-bodies or stress granules, respectively. (Scale bars = 10 μm)

Flow cytometry data was further confirmed by fluorescence microscopy analysis. 293T cells were transfected with Broccoli-plasmids and 24 hours later incubated with DFHBI and analyzed under a microscope. As observed by flow cytometry, both the number and intensity of mCherry-positive cells decreased with increasing copy number of the aptamer, and Broccoli intensity did not increase above 16× (**Fig. 2A**). While 4×, 8× and 16× Broccoli displayed uniform cytoplasmic staining, >32× resulted in the formation of green fluorescent aggregates. From 4× to 32× Broccoli, green fluorescence was mainly cytoplasmic, indicative of efficient mRNA export. For the 64× and 128× Broccoli constructs, green fluorescence accumulated in the nucleus, indicative of impaired mRNA export. This was more prominent in NIH-3T3, MEF and HeLa cells than in 293T (data not shown).

Broccoli analysis by microscopy represented an additional challenge, as fluorescence was very weak and bleaching occurred rapidly even when using highly sensitive electron-multiplying EMCCD cameras and low 488 nm laser excitation (**video 1.1** and **2.1**) at higher frame rates. However, Broccoli, albeit exhibiting higher stability than its predecessors, still is a quite unstable aptamer [3]. As such it was possible to do time-lapse imaging of cells at low frame rate such that fluorescence could recover by fluorochrome exchange in a PAINT-like fashion [11] (video **1.2** and **2.2**) but without single-molecule sensitivity.

Due to the highly structured and repetitive nature of the Broccoli concatemers, we hypothesized that 64× and 128× Broccoli would induce RNA aggregation. This might result in p-body or stress-granule formation, which are composed of arrested mRNAs or non-translating RNAs and their associated RNA-binding proteins [12], [13]. The respective Broccoli constructs might thus provide a versatile tool to study these interesting cellular structures. To test this hypothesis, we transfected 293T cells with Broccoli plasmids and co-transfected with an additional plasmid expressing either DCP1a-SNAP or DYRK3-SNAP, for the detection of p-bodies or stress-granules, respectively. 24 hours post-transfection, we incubated the cells with DFHBI and Alexa-647-SiR, to label SNAP, and performed live cell imaging. We found that transfection with Broccoli plasmids induced the formation of p-bodies, as shown by the presence of DCP1a positive granules (**Fig. 2B**, top two panels, cyan color) but not stress granules (**Fig. 2B**, bottom two panels, cyan color). Interestingly though, Broccoli aggregates were seen nearby but not co-localized to DCP1a aggregates. We conclude that Broccoli aggregates do not correspond to p-bodies or stress-granule sites.

To check the integrity and quantify mRNAs expressed from our Broccoli reporters, we isolated total RNA from transfected 293T cells and performed Northern blots. Both mCherry and Broccoli sequences were independently probed for. In accordance with the immunofluorescence data shown in Figure 1 and Figure 2A, both mCherry and Broccoli signals decreased with increasing copy numbers of Broccoli (**Fig. 3A** and **3B**, respectively). Both mCherry and Broccoli signals fell below the detection limit from 64× Broccoli onwards indicative of impaired mRNA stability.

**Figure 3.**
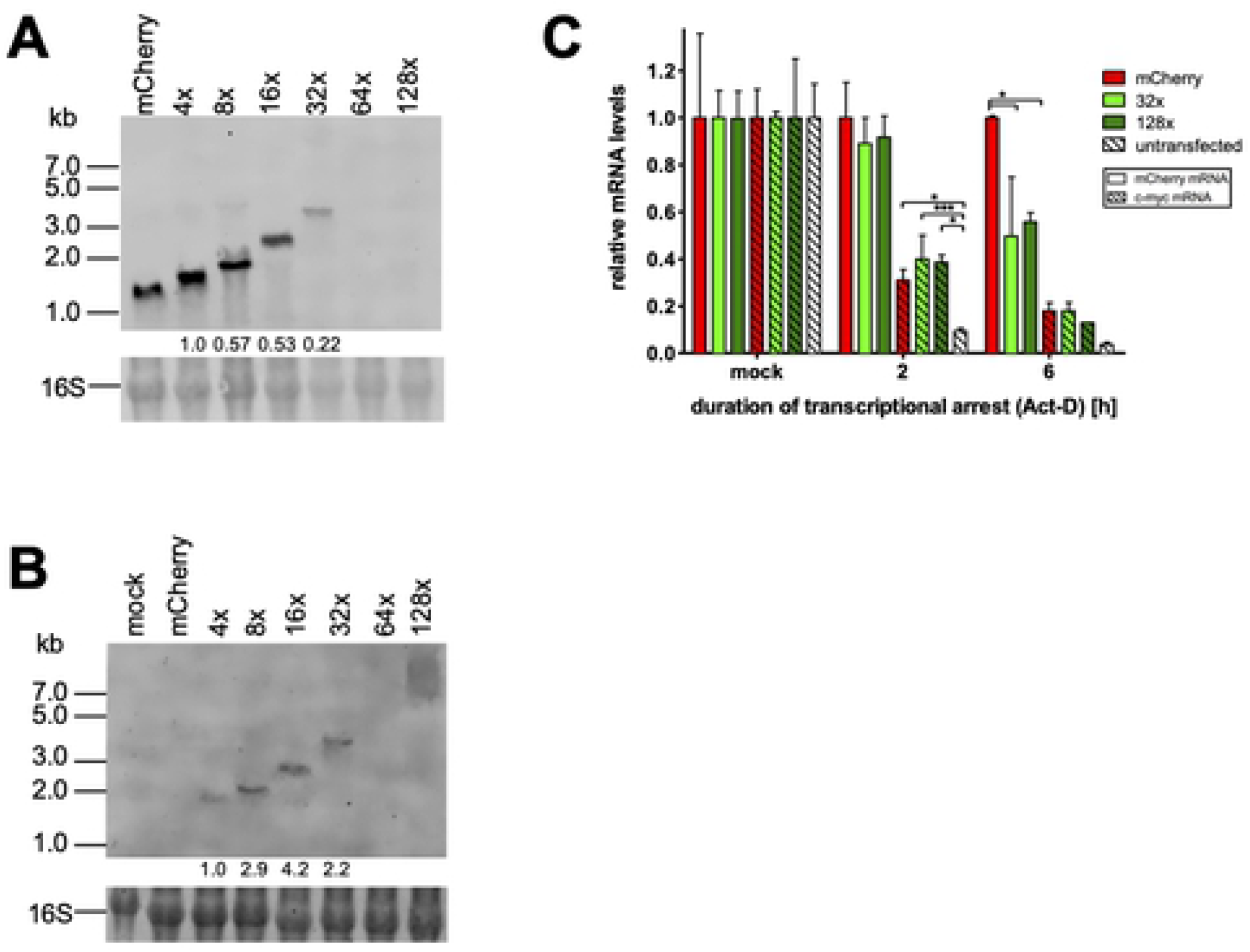
RNA of Broccoli is detectable up to 32 copies by Northern Blot. 293T cells were transfected with mCherry or Broccoli plasmids and 10μg of isolated total RNA loaded on a denaturing gel for Northern blot. A fluorescently-labelled probe against mCherry (a) or Broccoli was used to detect the respective RNAs. Intensities of mCherry and Broccoli RNA bands were measured using arbitrary values and normalized to either mCherry (a) or 4xBroccoli (b). (c) 293T cells were transfected with mCherry, 32× and 128× Broccoli plasmids. 24hours post-transfection cells were collected (mock) or treated with actinomycin D for 2 or 6 hours. RNA was isolated at each time point and RT-qPCR performed for detection of mCherry, c-myc and GAPDH. Empty bars show mCherry transcription levels, striped bars represent c-myc transcription levels. Statistical analysis performed by 2-way ANOVA. * P<0.05, **P<0.01, *** P<0.001 and mean fluorescence intensity of mCherry (d). (c) The values shown were normalized to the cells transfected with 4xBroccoli plasmid. Statistical analysis performed by t-test (c, d). * P<0.05, **P<0.01

To directly confirm the effect of Broccoli concatemers on mRNA stability, we employed the RNA polymerase inhibitor Actinomycin D (Act-D) and performed pulse-chase experiments. 24h after transfection of 293T cells with mCherry only, 32× or 128× Broccoli plasmids, we applied Act-D for 2 and 6 h. Cells were harvested at the respective time points and qRT-PCR was performed for mCherry. The highly unstable c-myc mRNA and the house-keeping gene GAPDH were used as a control. As expected, c-myc detection levels progressively decreased from 2 to 6 h post Act-D treatment (**Fig. 3C**). Interestingly, c-myc reduction was less pronounced in transfected cells regardless of the plasmid content, indicating that a transfection-induced stress response may interfere with cellular mRNA decay pathways. While mCherry mRNA levels dropped about 2-fold faster in presence of 32× Broccoli consistent with impaired mRNA stability, no further decrease in stability was observed for 128× Broccoli. In contrast to the highly unstable c-myc mRNA, the mRNA half-life of both the 32× and 128× Broccoli mRNA was in the range of about 6 h. While decreased mRNA stability thus accounts for the differences between the mCherry only and 32× Broccoli mRNA, it does not explain the additional loss in both mCherry and Broccoli Northern blot signal observed for the 64× and 128× reporters.

### Broccoli is not detectable by fluorescence in stable cell lines

The study of transcription and protein expression from plasmids needs to consider features such as transient expression of genes; transfection efficiency between different plasmids; and variation in copy number of plasmids per cell. To circumvent these variables, we generated polyclonal 293T and HeLa cell lines that express either mCherry or 16xBroccoli. Cell lines, expressing higher copy number of Broccoli were not possible to generate in our lab, as multiple attempts never resulted in full length Broccoli insertions but only recovered Broccoli fragments. No Broccoli fluorescence was detectable in the respective cell lines containing 16xBroccoli by FACS or fluorescence microscopy (**Fig. 4A**). Nevertheless, the 16xBroccoli cells showed reduced fluorescence of mCherry compared to the mCherry only cells consistent with the Broccoli-induced impairment of mRNA stability. Accordingly, Northern blot analysis from total RNA isolated from the cell lines revealed lower levels of mCherry RNA in 16xBroccoli-cells (**Fig. 4B**). In 16xBroccoli-HeLa cells, the RNA of mCherry did not reach detection levels. Furthermore, mCherry detection by RT-qPCR revealed an approximate 80% drop in RNA levels in 16xBroccoli-cells (**Fig. 4C**). To assess if we successfully integrated the full length of the 16 copies of Broccoli into our cells, we isolated genomic DNA and run a standard PCR using a forward primer at the 3’end of the mCherry sequence and a reverse primer at the 3’end of the final Broccoli repeat, covering the full length of Broccoli (where the expected length for 16xBroccoli would be 1278 bp). As a control, we performed PCR with the same primers on the lentiviral vector used to generate the cell lines. As expected, in the mCherry control vector and mCherry-cells we were unable to detect a Broccoli PCR product (**Fig. 4D**). When the 16xBroccoli-vector was included as template we detected a band with the expected size of the full-length 16xBroccoli (**Fig. 4D**). However, the band detected in both the 293T and HeLa cells expressing Broccoli were of smaller size, suggesting that the cells internally recombined some of the 16 copies of Broccoli sequence and generated a truncated sequence of Broccoli repeats. Nevertheless, the length of the product indicated that at least 8× copies of Broccoli had been maintained. (**Fig. 4D**).

**Figure 4.**
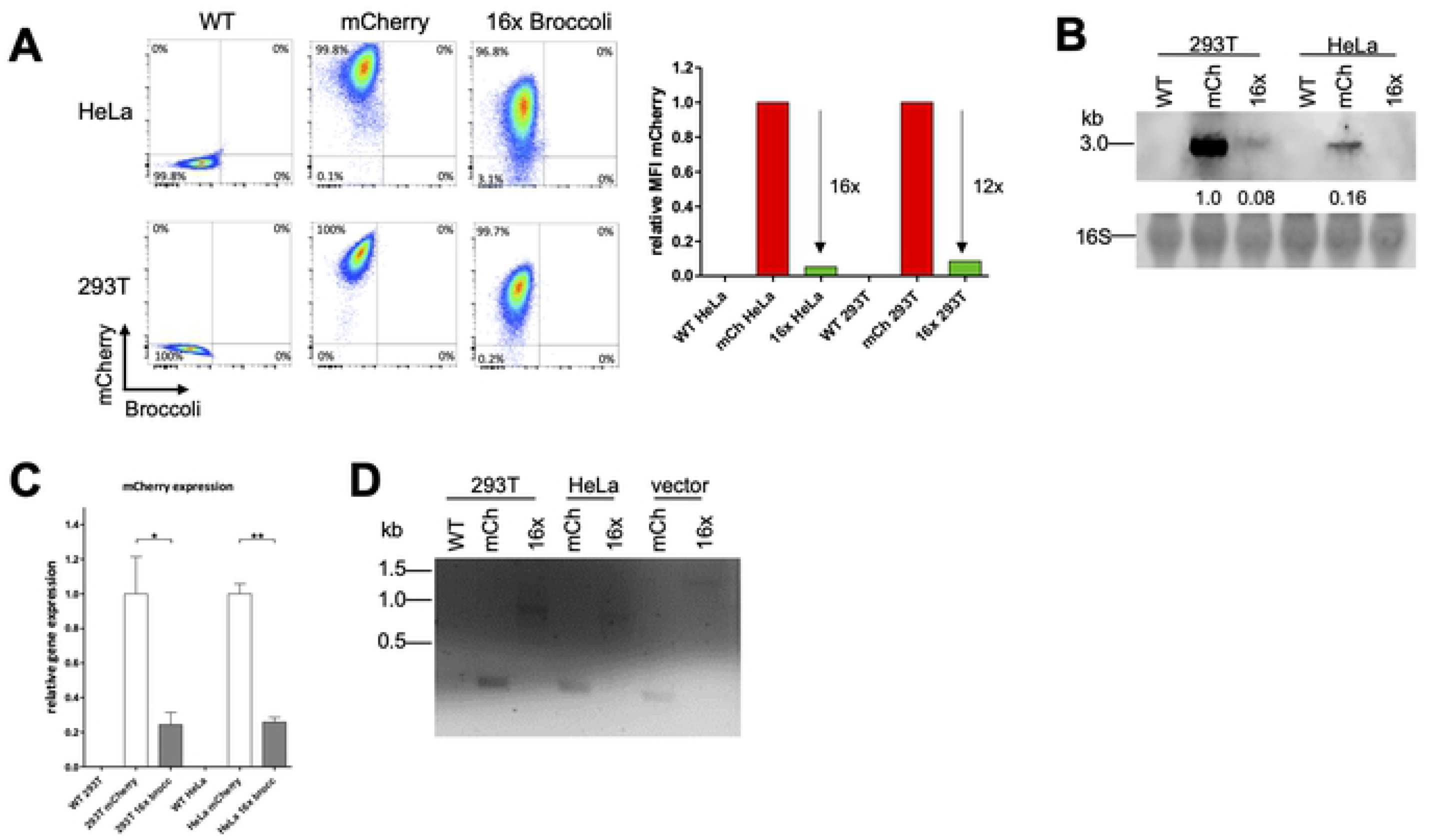
Polyclonal Broccoli-expressing cell lines showed reduced mCherry expression and no detectable Broccoli fluorescence. (a) Flow cytometry measurement of mCherry intensity in WT, mCherry or 16xBroccoli 293T or HeLa cells. (b) Northern Blot of mCherry RNA in the stable cell lines. RNA intensity was quantified and calculated relative to the 16S ribosomal band. All quantifications were further compared to the mCherry RNA in the mCherry-293T cells which was normalized to one. (c) RT-qPCR of mCherry. RNA was isolated from the stable cell lines, cDNA prepared and mCherry quantified by qPCR. Delta delta Ct of mCherry was calculated using GAPDH as the reference gene. Afterwards, all values were normalized to mCherry 293T cells. (d) Standard PCR for Broccoli from DNA isolated from 293T and HeLa cells expressing mCherry or 16× Broccoli (four lanes on the left side); lentiviral plasmid used for the generation of the mCherry and 16× Broccoli cell lines (two lanes on the right side). Statistical analysis performed by t-test (c). * P<0.05, **P<0.01

### Broccoli is not sustained by the genome of mouse cytomegalovirus

RNA aptamers represent interesting tools to visualize viral gene expression. Cytomegalovirus genomes are easily manipulated using reverse genetics approaches [14]. The large genome of murine cytomegalovirus (MCMV) provides an interesting vector to insert long DNA sequences such as Broccoli to study viral gene expression at the single-cell level. In order to reduce the risk for homologous recombination and thus removal of the inserted Broccoli concatemers, we took advantage of a large deletion mutant that lacks the first 17 genes of MCMV (delm01-m17) but shows wild-type virus replication properties in fibroblasts *in vitro* [15]. We inserted either mCherry or its 16× and 32× Broccoli variants using traceless mutagenesis [14] driven by a CMV promoter. Correct insertion and integrity of the MCMV BAC containing 16× and 32× Broccoli was confirmed by restriction digests. We subsequently reconstituted the respective viruses expressing either mCherry only (MCMV-mCherry), 16xBroccoli (MCMV-16x) or 32xBroccoli (MCMV-32x). We infected NIH-3T3 cells directly from the reconstituted virus inoculum. At 6h p.i., we already observed an average of 80% infection rate in the samples infected with MCMV, which was evaluated by analyzing the percentage of mCherry positive cells (**Fig. 5A**). At 2, 4 and 6 h p.i. a progressive increase in the mCherry mean fluorescence intensity (MFI) was evident (**Fig. 5B**). As expected, cells infected with the MCMV-16× and MCMV-32× displayed lower mCherry levels than MCMV-mCherry only infected cells (**Fig. 5B**). However, no Broccoli signal could be observed above background (**Fig. 5A, B**). To check the integrity of the inserted Broccoli concatemers, we isolated DNA from cells infected with MCMV-mCherry, MCMV-16× and MCMV-32×, at 6 hours post-infection. We designed primers that bind at the 3’end of mCherry and at the 3’end of Broccoli in order to obtain the full length of the Broccoli concatemers by PCR (expected length for MCMV-16× is 1200 bp, and MCMV-32× is 2292 bp). For MCMV-16×, we detected one product of the correct size, however the PCR also yielded three more bands below 1000 bp indicative of genomic recombination (**Fig. 5C**, middle lane). For the MCMV-32× sample, we did not observe a band matching the full length of Broccoli (**Fig. 5C**, right lane). Instead, we detected three products at approximately 800, 600 and 400 bp. We conclude that concatemers of Broccoli are not sufficiently stable to be faithfully maintained within the MCMV genome even for a very limited number of passages.

**Figure 5.**
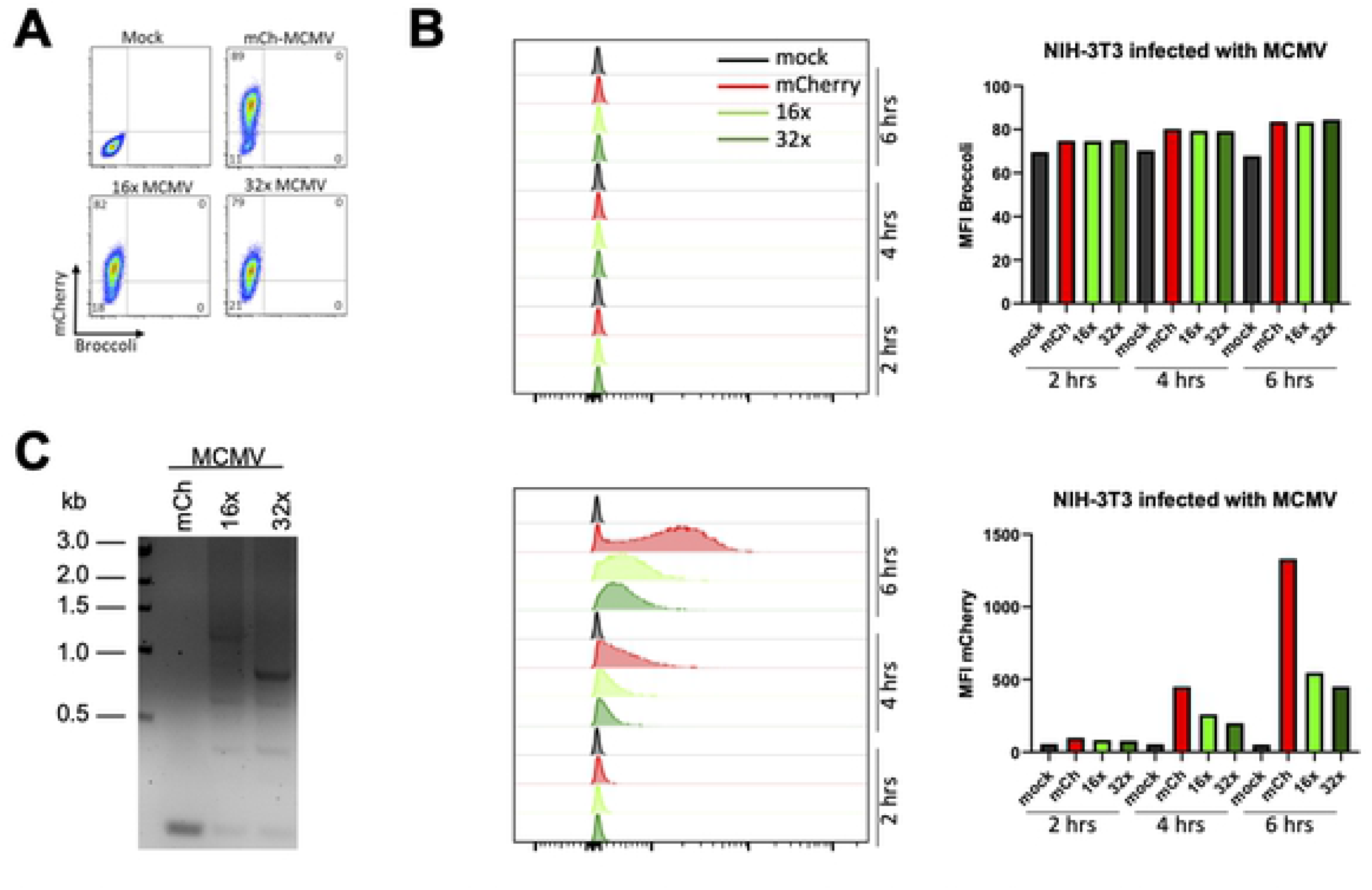
NIH-3T3 cells infected with Broccoli-MCMV viruses show reduced mCherry expression but no detectable Broccoli fluorescence. (a, b) NIH-3T3 cells were infected with undetermined MOI of mCherry-MCMV, 16x-MCMV or 32x-MCMV and analyzed by flow cytometry at 2, 4 or 6 hours post-infection. (a) Dot plots show cells at 6 hours post-infection. (b) Histograms and graph show progression of the viral infection/replication as shown by the mean fluorescence intensity of mCherry. (c) Standard PCR for Broccoli from DNA isolated from NIH-3T3 cells infected with mCh-MCMV, 16x-MCMV and 32x-MCMV, at 6 hours post-infection.

### Insertion of exonic sequences improves Broccoli fluorescence

Apparently, Broccoli concatemerization beyond 8 copies resulted in RNA aggregation and impaired both RNA localization and stability. In order to improve folding capacity of Broccoli, we created “spacers” between each 8 copies of Broccoli by insertion of partial sequences from exon 3 (151 nucleotides) or exon 5 (140 nucleotides), from the human GAPDH gene, at the 5’ end of Broccoli (**Fig. 6A**). Six new Broccoli expression plasmids were generated. We then transfected 293T cells and analyzed Broccoli and mCherry fluorescence by flow cytometry. While the percentage of transfected cells was similar between the exon-inserted and parental plasmids (**Fig. 6B**), the insertions resulted in an increase in the MFI of Broccoli for both the exon5- and exon3- inserted plasmids (**Fig. 6C**). Cells transfected with exon3-inserted plasmids showed significantly higher Broccoli MFI compared to exon5-inserted plasmids. This data suggests that Broccoli signals can be improved by optimizing the concatemers. Interestingly, the insertion of exons in between Broccoli repeats improved the MFI of mCherry, especially in cells transfected with 16xex3Broccoli compared to 16xBroccoli and in cells transfected with both 32xex3 and 32xex5 Broccoli compared to 32xBroccoli.

**Figure 6.**
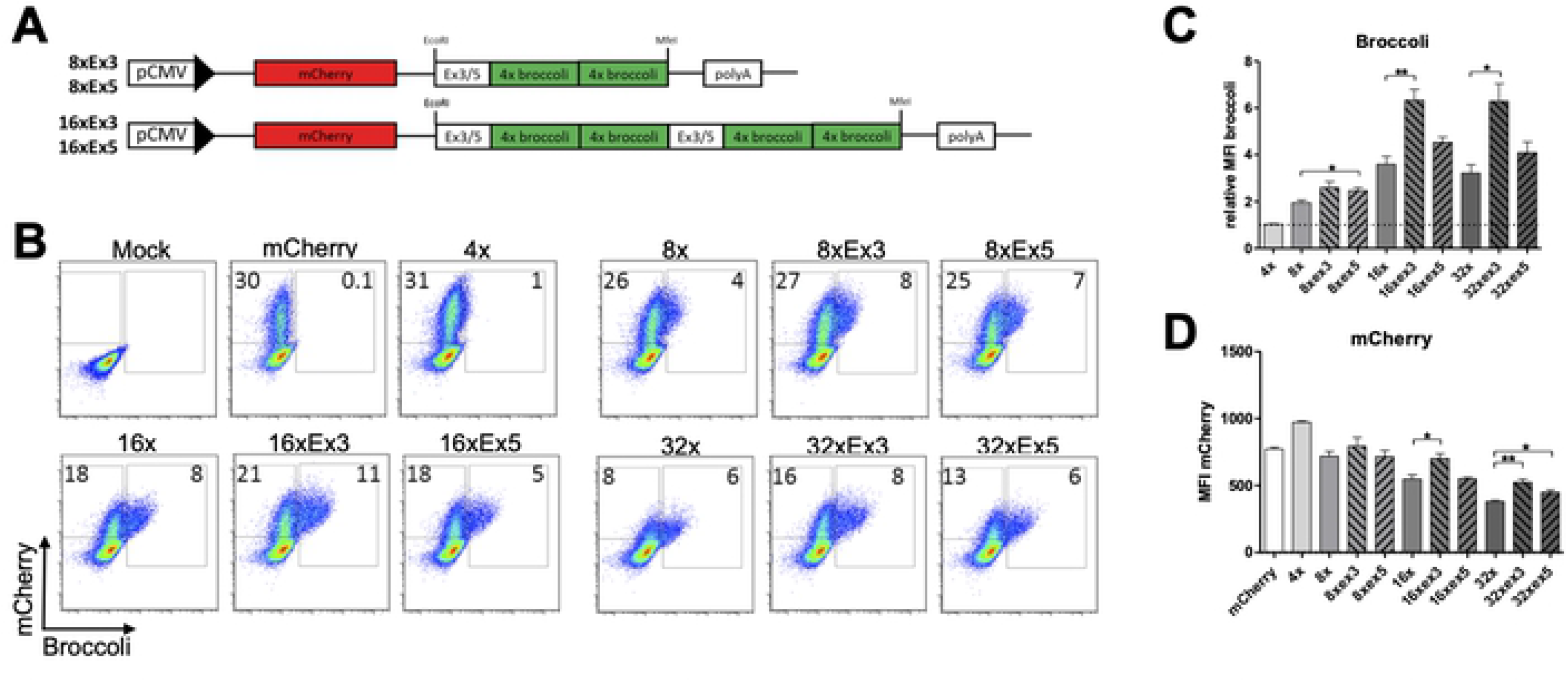
Expression profile of Broccoli tandem constructs with exonic inserts. (a) Cloning schematic of Broccoli with inserted exonic sequences, exon 3 or exon 5, from GAPDH gene of NIH-3T3 cells. (b, c, d) Flow cytometry data of 293T cells transfected with the different plasmids, collected and analysed 24 hours post-transfection. Comparison between the plasmids in regard to percentage of Broccoli and mCherry positive cells (b), mean fluorescence intensity of Broccoli

## Discussion

Fluorogenic RNA aptamers have the potential of surpassing the state-of-the-art real-time RNA imaging technology. They might induce less unexpected effects than fluorogenic proteins tagged to RNA, thereby delivering pictures of more physiological RNA transport and dynamics. Here we report on the development of such an approach, tandem-Broccoli arrays, with the scope of utilizing the constructs in imaging mRNA transport and dynamics in herpes virus infection.

We found that fluorescence signals of Broccoli concatemers increased linearly up to eight Broccoli repeats within an mCherry-expressing mRNA. However, no further increase in signal could be achieved with larger concatemers, over 16 repeats. Moreover, further extension of the concatemers resulted in a loss of Broccoli signal, RNA aggregation, impaired nuclear export, reduced RNA stability and impaired mCherry expression. Interestingly, we observed the formation of aggregated green-fluorescent RNA “granules” in the cytoplasm. Transfection of high-copy Broccoli concatemers triggered the formation of p-bodies but not of stress granules. Interestingly, the Broccoli aggregates did not colocalize with the induced p-bodies. Considering, the potential of Broccoli-RNAs to also form intermolecular interactions based on sequence complementarity, high-copy Broccoli concatemers may trigger liquid-liquid phase separation, not unlike of p-body formation [12]. Such RNA interaction-phase separation is a known mechanism that plays a role in disease developments [16]. As Broccoli aggregates did not localize to p-bodies, and neither to stress granules, we hypothesize that the presence of high amounts of this highly structured RNA induces a cellular stress response even though Broccoli RNAs are not recruited to these particular structures.

Aggregation of Broccoli-containing mRNAs involves intermolecular interactions between multiple Broccoli-carrying RNAs, which are by default complementary to each other. Interestingly, the insertion of linker sequences between every eight Broccoli repeats improved both Broccoli and mCherry signals. The introduction of a sequence from GAPDH exon 3, predicted to exhibit low secondary structures, led to a strong boost of Broccoli signal, actually surpassing the theoretical doubling of fluorescence intensity. This illustrates that the availability of sufficient space for Broccoli folding may be an important prerequisite for obtaining high signal Broccoli reporters. Nevertheless, we were still unable to increase fluorescence above the level provided by 16× Broccoli repeats.

Undesired Broccoli aggregation and plateauing of signal has not been reported by previous publications utilizing similar approaches though. In bacteria, Broccoli’s predecessor Spinach continues increasing fluorescence, albeit only reaching ~17-fold enhancement by stringing 64 repeats together [9]. Here the inducibility, and not general overexpression, by a lac operon might reduce unwished behavior. The most successful approach so far in this context though was carried out by Li *et al.* utilizing a newly designed substrate, termed BI, that increases Broccoli folding and stability.

We also set out to establish cell lines which stably express Broccoli. However, we were unable to generate cells expressing high copy numbers of Broccoli using lentiviral vectors. The same was true when expressing Broccoli concatemers from the murine cytomegalovirus genome. Recombination occurred despite the MCMV genome being undersized due to the deletion of the first 17 MCMV genes. We conclude that the expression of Broccoli concatemers negatively impacts on productive virus infection presumably due to the associated stress response, resulting in at least partial elimination of the Broccoli repeats within a few virus passages.

Finally, we set out to find a way of utilizing the Broccoli aptamer in real-time imaging. This was complicated by the rapid bleaching of signal that occurs when exciting the fluorophore bound by RNA. However, the bleached signal recovered over time, presumably due to the dissociation of bleached substrate from the RNA scaffold and binding of new DFHBI or its derivatives. Using sufficiently long-time intervals between excitation should therefore enable long-term time-lapse imaging of RNA abundance.

Overall, we show here that RNA aptamer concatemers can be rapidly manipulated to enhance real-time RNA tracing, but strategies need to be developed and fine-tuned to overcome their, impact on key aspects of RNA biology and avoid their rapid removal by homologous recombination.

## Material and methods

### Broccoli plasmids

We took advantage of the commercially available F30-2xdBroccoli construct (Geneart, Thermo Fisher) which is 234 nucleotides long. Short stem loop sequences to separate Broccoli aptamers from each other were added to this construct. The final F30-2xdBroccoli construct with the stem loops was flanked with EcoRI and MfeI sites, which were used to manifold the sequence to generate Broccoli concatemers. A single module, between EcoRI and MfeI, consists of 273 nucleotides and four DFHBI binding sites. This sequence was synthesized by Geneart (Thermo Fisher). We used a pCMV vector expressing mCherry downstream of a CMV promoter, and inserted Broccoli between the stop-codon of the mCherry protein and the SV40 poly-A-signal. To generate different-length Broccoli concatemers (**Table 1**), the Broccoli sequence was extracted using EcoRI and MfeI and reinserted into the linearized vector, cut only with EcoRI, effectively doubling the sequence with each iteration (**Fig. 1**).

To overcome folding restrictions caused by the length of the Broccoli constructs, we inserted exonic sequences from the GAPDH gene of NIH-3T3 cells, at the 5’end (EcoRI site) of 8 copies (8x) Broccoli. These exonic sequences (see supplementary information for exact sequence) were PCR amplified, with primers containing EcoRI or MfeI sites (prW166 and prW167 or prW168 and prW169, **Table 1**), from exon 3 (151 bp length) or exon 5 (140 bp length) of the gene. The longer Broccoli concatemers containing exons were synthesized following the same cloning strategy described above (**Fig. 6**).

### DCP1a-SNAP and DYRK3-SNAP plasmids

pT7-EGFP-C1-HsDCP1a was a gift from Elisa Izaurralde (Adgene plasmid #25030) [17] and pDONR223-DYRK3 was a gift from William Hahn and David Root (Adgene plasmid #23635) [18]. The SNAP ICAM-1 fusion protein was inserted before both the DCP1a and DYRK3 sequences for further use in fluorescence microscopy and detection of p-bodies and stress-granules, respectively. Therefore, the SNAP sequence was first amplified by PCR using the primers prW520 (**Table 2**) containing an AgeI site and prW521 containing a BspEI site, and inserted into the DCP1a plasmid by replacing the EGFP sequence. We will refer to this plasmid as DCP1a-SNAP. Afterwards, we amplified the DYRK3 sequence from the purchased plasmid using the primers prW358 and prW359 containing 15bp complementary sites to DCP1a plasmid. DYRK3 sequence was inserted using the In-Fusion HD cloning kit (Takara Bio USA, Inc), to replace the DCP1a sequence with DYRK3. This plasmid will be referred to as DYRK3-SNAP.

### Cell lines

NIH-3T3 cells were grown in Dulbecco’s modified eagle medium (DMEM, Gibco) containing 10% newborn calf serum (NCS) and penicillin-streptomycin. HEK-293T and HeLa cells were grown in DMEM containing 10% fetal bovine serum (FBS) and penicillin-streptomycin. M2-10B4 cells were grown in RPMI containing 10% FBS and penicillin-streptomycin.

### Generation of the HeLa and HEK-293T cell lines expressing mCherry or mCherry-16× Broccoli

The pHRSIN.pSFFV MCS(+) pGK Hygro plasmid (referred from here onwards as pHRSIN) was kindly provided by Paul Lehner. MCherry or 16 copies (16x) of Broccoli was inserted into the multicloning site of the pHRSIN plasmid. The pHRSIN plasmid was digested with BamHI, followed by Klenow (New England Biolabs) treatment and afterwards KpnI and CIP. mCherry and 16xBroccoli plasmids were digested with AfeI and KpnI. Ligation of the correct sequences followed, using T4 ligase and incubation at room temperature. Afterwards, MAX Efficiency Stbl2 competent cells (Invitrogen) were transformed by heat-shock with the ligation reaction and grown in LB agar plates for clone selection. For confirmation, clones with the correct insertions were sequenced before proceeding. Due to the length of all constructs above 16xBroccoli, further cloning with the remaining Broccoli repeats into pHRSIN was not successful. HEK-293T or HeLa cells were transfected with pHRSIN, pHRSIN-mCherry, or pHRSIN-16xBroccoli plus psPAX2 and CMV.VSVg plasmids at a ratio 5:3:1, respectively. Following transfection, we collected the supernatant 48hrs later and transduced new HEK293T or HeLa cells. 48 hours later, cells were split and grown in media supplemented with 150μg/ml hygromycin for positive selection.

### Broccoli expression and localization

HEK-293T cells were transfected with 4×, 8×, 16×, 32×, 64× or 128× Broccoli plasmids or a mCherry plasmid (control) in equimolar ratios, using the transfection reagent polyethylenimine (PEI, Polysciences). (Note: due to large differences in length between plasmids (see Table 1 for base pair number information), we co-transfected with a control plasmid, pEPkan-S (Adgene plasmid #41017) [19], which is not containing a sense-ORF for these cells, to equalize the DNA mass of each transfection sample). 24 hours later, the cell media was aspirated and PBS with 20μM GK132 DFHBI (more stable) or DFHBI-1T (less stable) was added and cells incubated at 37°C for 30 minutes. Broccoli was either analyzed by flow cytometry (LSRII, BDbioscience) using DFHBI-1T or microscopy (GK132 DFHBI). HEK-293T and HeLa cell lines generated to express mCherry or 16× Broccoli were trypsinized, washed with PBS, incubated with PBS plus 20μM of DFHBI-1T for 30 minutes, and signals acquired by flow cytometry.

To assess transfection efficiency of the different Broccoli plasmids, HEK-293T cells were transfected with mCherry, 32× or 128× Broccoli together with 25ng of the plasmid pEGFP-C1 (adgene plasmid #6084-1). Cells were collected 24 hours post-transfection and analyzed by flow cytometry for mCherry versus GFP expression (no DFHBI was added).

**Table 1:**
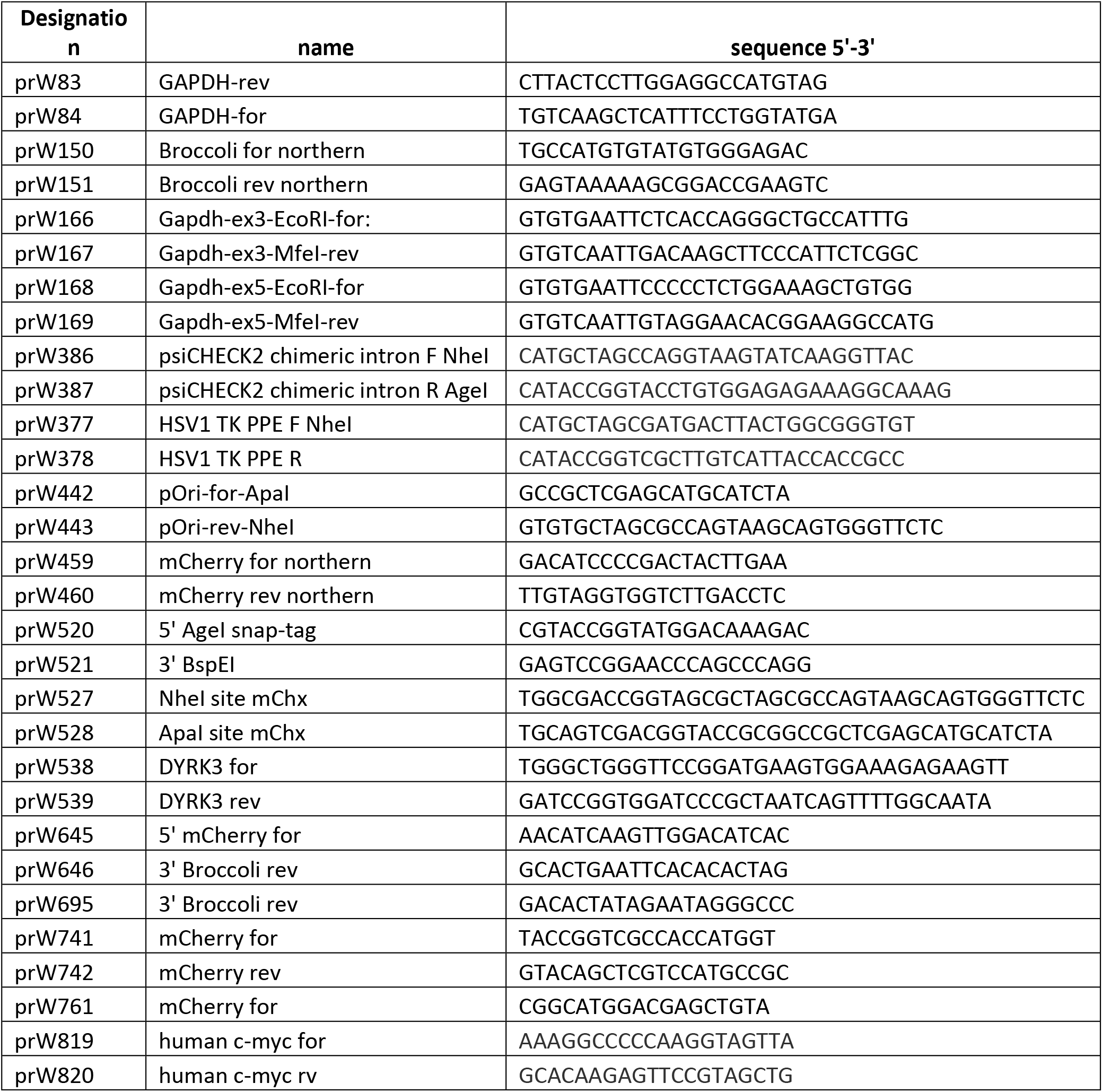
Primers

### P-bodies and stress granules co-localization

HEK-293T cells were co-transfected with either 16×, 32×, 64× or 128× Broccoli plasmids together with DCP1a or DYRK-SNAP. Next day, the cells were labelled with Hoechst, PFP-DFHBI for Broccoli and SNAP-Cell 647-SiR for SNAP and imaged using a Nikon spinning disc system consisting of a Yokogawa W2 and two Andor iXON888 cameras using NIS-Elements for image acquisition. A Nikon 100× 1.49 NA Apo-TIRF objective was used resulting in 130nm pixel size. The system was equipped with standard 405, 488, 561, 640 nm laser lines and corresponding filter sets. Since Broccoli-fluorescence was weak and sometimes barely distinguishable from cellular autofluorescence, Broccoli-expressing cells were identified in the 488-channel through their rapid bleach-behavior in live-camera mode. Upon identification, excitation was paused for 30 seconds until Broccoli fluorescence recovered and a single 4-channel Image (Hoechst, Broccoli, mCherry, SNAP) was taken using Nikon NIS AR 4.5 software.

### Northern Blot

HEK-293T cells were seeded on 10-cm dishes and transfected using PEI with 25μg of DNA at a ratio 1:3 DNA (mass): PEI (volume). 24 hours post-transfection, supernatant was aspirated and 2mL of TRI Reagent (Sigma Aldrich) were added per 10-cm dish to resuspend cells to a homogeneous lysate. RNA was purified as follows: 600μL of chloroform were added to each lysate, samples were vortexed vigorously and centrifuged; the aqueous phase was collected into a fresh tube where 1 volume of isopropanol was added and the samples were centrifuged; supernatants were removed and the precipitate washed two times with 75% ethanol and resuspended in 50μl nuclease free water. To avoid DNA contamination, samples were submitted to DNase treatment using the Turbo DNA-free kit (Life Technologies) and the “rigorous DNase treatment” protocol according to the manufacturer’s instructions. RNA was further purified by adding 1 volume of Roti-Aqua-P/C/I (Roth) followed by two 75% ethanol washes as described above. 10μg (for mCherry detection) or 20μg (for Broccoli detection) of RNA was resolved on a 1% agarose denaturing gel followed by blotting on a nylon membrane (Amersham Hybond-N^+^, GE Healthcare). A probe for mCherry or for 4× Broccoli was synthesized by PCR using the 4× Broccoli plasmid as template and the primers prW150-prW151 for Broccoli and prW459-prW460 for mCherry. The probes were fluorescently labelled by including 10nM of Aminoallyl-dUTP-ATTO-680 (Jena Bioscience) in the PCR reaction. For hybridization, the membrane was incubated overnight at 60°C together with the probes diluted in Church buffer (1% (w/v) BSA, 1mM EDTA, M phosphate buffer, 7% (w/v) SDS). The probes were detected via the 700nm channel of the LI-COR Odyssey Imaging System.

### Broccoli insertion into mouse cytomegalovirus

Insertion of Broccoli fragments into the BAC C3× delm01-m17 mutant (here onwards referred as pD2-BAC, previously described [15] and a kind gift from Zsolt Ruzsics), was performed using the FLPe-FRT system as follows. A transiently expressed pOri plasmid containing two Frt sites was linearized using ApaI. Afterwards, a NheI site was inserted by PCR-mutagenesis and amplification of the vector using primers prW442 (containing an ApaI site) and prW443 (containing a NheI site). The derived PCR product (pOri plasmid with ApaI and NheI sites) was used to ligate with mCherry or Broccoli fragments digested ApaI and NheI as well, resulting in the plasmids pOri-mCherry; pOri-16xBroccoli and pOri-32xBroccoli. Electro-competent E.Coli (GS1783), containing the pD2 BAC, were transfected with the different pOri plasmids together with a helper plasmid (pGPS-FLPe, a kind gift from Zsolt Ruzsics) which expresses the FLPe-recombinase. This led to the generation of different C3× BAC hereon referred to as MCMV-mCherry, MCMV-16× and MCMV-32× (**Table 2**).

**Table 2:** Constructs in plasmids

**Constructs in plasmids (Table 2):**
| Name | Description | mCherry-Broccoli Size (bp) |
| --- | --- | --- |
| mCherry | mCherry | 711 |
| 4xBroccoli | mCherry-4xBroccoli | 1005 |
| 8xBroccoli | mCherry-8xBroccoli | 1278 |
| 16xBroccoli | mCherry-16xBroccoli | 1824 |
| 32xBroccoli | mCherry-32xBroccoli | 2916 |
| 64xBroccoli | mCherry-64xBroccoli | 5100 |
| 128xBroccoli | mCherry-128xBroccoli | 9468 |
| 8xex3 Broccoli | mCherry-8xBroccoli-exon3 GAPDH | 1435 |
| 16xex3 Broccoli | mCherry-16xBroccoli-exon3 GAPDH | 2138 |
| 32xex3 Broccoli | mCherry-32xBroccoli-exon3 GAPDH | 3544 |
| 64xex3 Broccoli | mCherry-64xBroccoli-exon3 GAPDH | 6356 |
| 8xex5 Broccoli | mCherry-8xBroccoli-exon5 GAPDH | 1424 |
| 16xex5 Broccoli | mCherry-16xBroccoli-exon5 GAPDH | 2116 |
| 32xex5 Broccoli | mCherry-32xBroccoli-exon5 GAPDH | 3500 |
| 16x-intron-Broccoli | 16xBroccoli containing a chimeric intron | 1975 |
| 16x-TK-Broccoli | 16xBroccoli containing the HSV-1 thymidine kinase export signal | 1955 |
| 128x-intron-Broccoli | 128xBroccoli containing a chimeric intron | 9619 |
| 128x-TK-Broccoli | 128xBroccoli containing the HSV-1 thymidine kinase export signal | 9599 |
| MCMV-mCherry | pD2 CMV strain contain mCherry | 711 |
| MCMV-16x | pD2 CMV strain contain mCh-16xBroccoli | 1824 |
| MCMV-32x | pD2 CMV strain contain mCh-32xBroccoli | 2916 |

### PCR for Broccoli

To detect insertion of Broccoli sequences in the originated plasmids; cell lines; or MCMV, midipreps were prepared for each plasmid, and DNA was isolated from the designated stable cell line or from NIH-3T3 cells infected with MCMV. For plasmids 1-4ng were used for each PCR reaction, while for genomic DNA 50-100ng was used. The primer pair prW645-646 was used to detect Broccoli inserted in the lentivirus vector pHRSIN or in the stable cell lines, and the pair prW695-761 to detect Broccoli in MCMV. These primers flank the whole Broccoli sequence by annealing to the 3’ end of the mCherry sequence and the 3’ end of the last Broccoli repeat, respectively. The annealing temperatures used were: 61°C for prW645-646 and 50°C for prW695-761. Each PCR reaction mix included: Taq polymerase, 10× buffer, DMSO, 0.2mM of dATP, dTTP, dCTP, 0.05mM of dGTP (Roche) and 0.15mM of 7-deaza-dGTP (7-deaza-2′-deoxyguanosine 5′-triphosphate, NEB).

### Actinomycin D treatment and qPCR

HEK-293T cells were transformed with mCherry, 32× Broccoli or 128× Broccoli plasmids as described above. 24 hours post-transfection, 10μg/mL of actinomycin D was added to the cells. Three technical replicates were performed for each condition and plasmid. Cells were collected at 24hours post-transfection without any treatment (mock) or 2- and 6-hours post-treatment with actinomycin D. RNA was isolated from these cells and submitted to DNase treatment, as described above. From total RNA, 300ng were used to synthesize cDNA with the All-in-One cDNA synthesis supermix (Biotool) and according to the manufacturer’s instructions. Gene expression of mCherry was evaluated by a 2-step qPCR using 30ng of cDNA, the primers prW741 and prW742 and SYBR qPCR mastermix (Bimake). The primers prW83 and prW84 for GAPDH and prW819 and prW820 for c-myc were used as references/controls. SyBr green measurements were acquired using a Roche LightCycler 96 machine.

## Acknowledgments

This work was supported by Infect-ERA grant eDEVILLI to L.D. J.B. is funded by the Deutsche Forschungsgemeinschaft (DFG, German Research Foundation) under Germany’s Excellence Strategy – EXC 2155 “RESIST” – Project ID 390874280.

## Author contributions

Conceptualization: L.D., T.H., J.B.; Data curation: M.R., M.A.P.B; Formal analysis: M.R., M.A.P.B; Funding acquisition: J.B, L.D.; Investigation: M.R., M.A.P.B., A.W., J.B.; Methodology: M.R., M.A.P.B., L.D., T.H., N.W., J.S.; Project administration: L.D.; Resources: N.W., J.S.; Supervision: L.D.; Validation: M.R., L.D., J.B.; Visualization: M.R., M.A.P.B.; Writing – original draft: M.R., M.A.P.B., L.D., J.B.; Writing – review and editing: M.R., M.A.P.B, L.D., J.B.

## Supplementary information

To assess if transfection efficiency changes with the different plasmid sizes, we transfected 293T cells with mCherry, 32×, or 128× Broccoli (see detailed transfection procedure in the methods section) and co-transfected with a control eGFP-expressing plasmid. 24 hours post-transfection the cells were harvested and analyzed by flow cytometry (the excitation of Broccoli-DFHBI emits light in the same detector as GFP, therefore DFHBI was not added to these samples). Dot plots show that samples transfected with Broccoli plasmids have more GFP positive cells compared to the samples transfected with the mCherry plasmid (**Supplementary Fig. 1A**). Moreover, the overall fluorescence intensity of GFP is also higher in the Broccoli transfected samples (**Supplementary Fig 1B**). This data suggests that transcript length is a critical factor and here GFP is preferentially transcribed compared to mCherry where the mCherry gene is located at the 5’end of lengthy DNA sequences such as Broccoli.

To exclude the possibility that the differences in expression between mCherry and Broccoli, observed in the cells, were not the result of deteriorating quality of the plasmids, mCherry, 32xBroccoli and 128xBroccoli plasmids were prepared from three different bacterial cultures each, followed by plasmid purification, and resulting in three different batches for each plasmid type. Afterwards, each individual cell culture of 293T cells was transfected with a different batch of mCherry; 32xBroccoli plasmid; or 128xBroccoli. 24 hours post-transfection, cells were collected, total RNA isolated and RT-qPCR performed for mCherry detection, using GAPDH as reference. The results confirm a pronounced decrease in mCherry levels when cells are transfected with 32× and 128× Broccoli, excluding the hypothesis of plasmid deterioration (**Supplementary Fig. 1C**).

### Ex3-Insert sequence

5’-

tcaccagggctgccatttgcagtggcaaagtggagattgttgccatcaacgaccccttcattgacctcaactacatggtctacatgttccagtatgactccactcacggcaaattcaacggcacagtcaaggccgagaatgggaagcttgt-3’

### Ex5-Insert sequence

5’-

cccctctggaaagctgtggcgtgatggccgtggggctgcccagaacatcatccctgcatccactggtgctgccaaggctgtgggcaaggtcatcccagagctgaacgggaagctcactggcatggccttccgtgttccta -3’

**Supplementary Figure 1.** (a, b) 293T cells were co-transfected with a plasmid expressing eGFP plus mCherry, 32×, or 128xBroccoli plasmids. Percentage and intensity of eGFP and mCherry positive cells were detected by flow cytometry 24 hours later and are shown in the dot plots (a) and histograms (b), respectively. To accurately detect eGFP, DFHBI was not added to the cells. Three technical replicates are shown in the graphs. (c) The same protocol as in (d and e) was applied and cells were collected 24 hours post-transfection for RNA isolation. RT-qPCR was performed to measure mCherry expression. The values shown were normalized to mCherry transfected cells. Statistical analysis performed by t-test. * P<0.05, **P<0.01, *** P<0.001

**Supplementary Figure 2.** Expression profile of Broccoli tandem constructs with exonic inserts. (a) 293T cells were transfected with mCherry, 8xEx3, 8xEx5, 16xEx3, 16xEx5, 32xEx3, 32xEx5, 64xEx3 Broccoli plasmids and analysed by fluorescent microscopy. (b, c) Northern Blot from RNA isolated from 293T cells transfected with Broccoli plasmids containing exonic sequences. Northern membranes were probed for mCherry (b) or Broccoli (c). Values below the membranes and in the graphs indicate the intensity of the bands relative to the 16S ribosomal band for each sample and normalized to cells transfected with mCherry (b) or 4× Broccoli (c). Scale bar = 25μm

**Supplementary Figure 3.** Insertion of an HSV-1 intronic sequence or thymidine kinase did not change localization of nuclear 128× Broccoli. 293T cells were transfected with 16x-, 16x-intron-, 16x-TK-, 128x-, 128x-intron-, and 128x-TK-Broccoli plasmids and analyzed by microscopy (a) or flow cytometry (b, c). (b) Percentage of cells that express mCherry and Broccoli is shown in dot plots. (c) Comparison of Broccoli (top histogram and graph) and mCherry (bottom histogram and graph) expression between the different plasmids, as mean fluorescence intensity. Statistical analysis performed by t-test (c). * P<0.05, **P<0.01. Scale bar = 50μm

The chimeric intron was PCR amplified from psiCHECK-2 (Promega) using primers prW386 and prW387. The herpes simplex virus 1 strain Syn17+ thymidine kinase export signal was PCR amplified from infected cellular DNA following TRIreagent purification using primers prW377 and prW378. Amplicons were ligated into the NheI and AgeI sites (upstream of mCherry) of 16× and 128× Broccoli.

**Supplementary Figure 4.** Full images of blots and gels depicted in figures 3, 4 and 5.

### Supplementary Video 1.1

293T cells were transfected with mCherry 32× Broccoli and analysed by live cell spinning disk microscopy with consecutive 100 ms per frame. Rapid bleaching can be observed

### Supplementary Video 1.2

293T cells were transfected with mCherry 32× Broccoli and analysed by live cell spinning disk microscopy with 100 ms per frame every 5s. Apparent bleaching is mostly omitted likely due to dye exchange.

### Supplementary Video 2.1

293T cells were transfected with mCherry 32× Broccoli and analysed by live cell spinning disk microscopy with consecutive 100 ms per frame. Rapid bleaching can be observed

### Supplementary Video 2.1

293T cells were transfected with mCherry 32× Broccoli and analysed by live cell spinning disk microscopy with 100 ms per frame every 7s. Apparent bleaching omitted likely due to dye exchange.

